# Colon-to-hind paw cross-organ sensitization is partially mediated by TrkB.T1-facilitated spinal neuroinflammation to activate lumbar DRG neurons

**DOI:** 10.64898/2026.07.25.740685

**Authors:** Parshva Mehta, Namrata Tiwari, Cristina Smith, Shanwei Shen, Taggert Barton, Aron H Lichtman, Liya Y Qiao

**Author notes:** Corresponding Author: Liya Qiao, Ph.D., Department of Cellular, Molecular and Genetic Medicine Department of Neuroscience and Anatomy, School of Medicine Virginia Commonwealth University, Richmond, VA 23298-0551 USA. equal contribution (Parshva Mehta, Namrata Tiwari, Cristina Smith).

## Abstract

Patients with bowel disease can develop referred somatic pain at a later time with unknown molecular mechanisms. Using experimental mice with colitis induced by intracolonic installation of 2,4,6-Trinitrobenzenesulfonic acid (TNBS), we find an increase in the percentage of hind paw primary afferent neurons expressing Piezo2 or calcitonin gene-related peptide (CGRP), which are attenuated by TrkB.T1 knockout (KO). Concomitantly, TrkB.T1 KO also attenuates colitis-induced hind paw mechanical hypersensitivity and pain. Next, we find that TrkB.T1 is expressed in spinal cord astrocytes and its expression level is increased by colitis. TrkB.T1 KO reduces colitis-induced upregulation of Tumor necrosis factor-alpha (TNF-α) mRNA but not upregulation of interleukin (IL)-6 mRNA in the spinal cord. Using calcium (Ca^2+^) imaging and ex vivo approaches, we find that TNF-α elicits Ca^2+^ transients in capsaicin-sensitive as well as capsaicin-insensitive DRG neurons, and increases Piezo2 and CGRP expression in DRG neurons via distinct signaling pathways. Notably, TNFα-or colitis-induced Piezo2 upregulation in L4 DRG neurons is mediated by or associated with the PI3K/Akt pathway that does not participate in CGRP upregulation. In contrast, CGRP upregulation in hind paw primary afferent neurons is associated with an upregulation of phosphorylated cAMP response element binding protein (p-CREB). Finally, we find that TrkB.T1 KO does not change the expression level of transient receptor potential cation channel subfamily V member 1 (TrpV1) in L4 DRG in colitis, explaining the ineffectiveness of TrkB.T1 KO on colitis-induced hind paw thermal hyperalgesia. These results suggest complex and distinct molecular pathways in colitis-induced somatic pain modalities and provide information for specific pain modality management.

## INTRODUCTION

It is a common occurrence in human that pain is perceived at a remote location away from the injured organ, namely referred pain. The pain comorbidity causes a much greater rate of disability and psychological symptoms [1–3]. For example, heart attack can cause pain in the neck, shoulders, and back. Patients with bowel diseases also have back pain, leg pain and bladder disorders. This is also true in experimental rodents that induction of colonic inflammation results in hind paw mechanical hypersensitivity [4–7], manifested by an elevated calcium (Ca^2+^) level in L4 DRG [8] that contains primary afferent neurons of the hind paw [9]. Since primary afferent neurons that innervate the distal colon mainly locate in the thoracolumbar (T13-L2) and lumbosacral (L6-S1) DRGs [10, 11], it requires mediators to convey the aberrant sensory process from colitis-sensitized colonic afferent neurons to the L4 DRG, which is believed via the spinal central sensitization.

Neuropeptides released from DRG to the spinal cord play a critical role in spinal central sensitization. The well-defined such neuropeptides include brain-derived neurotrophic factor (BDNF) [12]. BDNF release to the spinal cord acts on its high affinity receptors, TrkB or TrkB.T1, to promote activation of neurons and astrocytes, respectively [13–17], thereby leading to spinal central sensitization. TrkB.T1 deletion from astrocytes attenuates nerve injury-induced neuropathic pain [17–20], suggesting a regulatory role of astrocytes in pain-sensing neuron activity. The TrkB and TrkB.T1 receptors share homologous N-terminal extracellular domain to have the same affinity to bind to BDNF, with a completely different C-terminus [21–23]. Unlike TrkB, TrkB.T1 does not possess intracellular tyrosine kinase activities but couples to Rho GTPase to activate the intracellular Ca^2+^ pathway, promoting vesicle trafficking and altering glial morphology [24, 25]. A clinical study in patients with irritable bowel syndrome (IBS) reveals that TrkB.T1 nucleotide polymorphisms is associated with worsened somatic symptoms [26], suggesting a role of TrkB.T1 in IBS-associated somatic dysfunction. However, the underlying molecular mechanism and pathways by which TrkB.T1 participates in colon-to-limb (e.g., hind paw in rodents) sensory crosstalk are not investigated.

One of the triggers of spinal central sensitization is malfunction of astrocytes in the spinal cord that alters the microenvironment, in the context of pain, generates neuroinflammation in pain induction and chronic maintenance [27–29]. The star-shaped mature astrocytes extend their processes to enwrap synapses in a ratio of 1 astrocyte to over 100k individual synapses to efficiently mediate neuron-glia-neuron crosstalk [30], support neuronal health and modulate neuronal activity [31–33]. Astrocytes also form a complex network with other astrocytes along the spinal cord therefore signals can be propagated along the cord [34]. In an animal model of persistent visceral pain caused by intracolonic installation of 2,4-dinitrobenzenesulfonic acid, a significant activation of astrocytes is observed in the dorsal and ventral horns of the spinal cord [35]. Proinflammatory factors produced by reactive astrocytes in the spinal cord in turn antidromically reinforce the activity of DRG neurons.

In DRG, Piezo2 is identified as a mechanical transducer to mediate mechanical sensation and pain [36–39]. Calcitonin gene-related peptide (CGRP) is also a potent pain marker that is positively associated with a number of pain modalities [40, 41]. The current study is undertaken to assess whether colitis-induced hind paw hypersensitivity is manifested by changes in the expression of Piezo2 and CGRP in L4 DRG and whether TrkB.T1 has a role in colon-to-hind paw cross-organ sensitization using experimental mice with colitis.

## MATERIALS AND METHODS

### Animals and breeding strategies

Animals used were C57BL/6J mice on which TrkB.T1 homozygous knockout (TrkB.T1^−/−^, or TrkB.T1 KO) was created and back-crossed, and generously provided by Dr. Lino Tessarollo [42, 43]. Age-matched male mice ((20–30 g, aging 8-12 weeks) were grouped for comparison. GCaMP6f mice were generated by crossing Ai95 mice (JAX Stock # 028865) with CMV-Cre mice (JAX Stock # 006054). Piezo2;mCitrine mice were generated by crossing Piezo2-EGFP-IRES-Cre mice (Piezo2-Cre, JAX Stock # 027719) [44] with R26-LSL-Gi-DREADD (JAX Stock # 026219). Standard husbandry conditions with 12:12-h light cycles and free access to regular food/water were supplied to each cage that housed 2–5 mice to ensure adequate social environment. All experimental protocols involving animal use were approved by Virginia Commonwealth University Institutional Animal Care and Use Committee (IACUC). Animal care was in accordance with the Association for Assessment and Accreditation of Laboratory Animal Care (AAALAC) guidelines.

### Labeling of hind paw primary afferent neurons by neuronal tracing dye

We injected neuronal tracing dye Fast Blue (FB, 4 %, weight/volume; Polysciences, Inc. Warrington, PA) into the plantar surface of the hind paw to back label hind paw primary afferent neurons in DRG. Specifically, under anesthesia (2.5 % isoflurane) and following sterilization, a total of 10 µL of FB was injected into 3-4 spots inside the glabrous skin of the hind paw plantar. Animals were allowed for recovery for a week and FB-labeled DRG neurons were assessed.

### Immunostaining

DRGs were fixed in 4% paraformaldehyde followed by incubation in 25 % sucrose overnight at 4 ℃ for cryoprotection. DRG sections (10 µm thickness) were processed for on-slide immunostaining. The primary antibodies were diluted in PBST (0.3 % Triton X-100 in 0.1 M PBS, pH 7.4) buffer containing 5 % normal donkey serum (Jackson ImmunoResearch, West Grove, PA). The primary antibodies used were rabbit anti-Piezo2 (1:500, Novus Biologicals LLC, Cat# NBP1-78624), rabbit anti-CGRP (1:1000, Invitrogen, Cat# PA5-114929), goat anti-CGRP (1:2000, Abcam, Cat# AB36001), rabbit anti-p-CREB (1:500, Cell Signaling, Cat# 9198), and rabbit anti-p-Akt (1:500, Cell Signaling, Cat# 4060), rabbit anti-TRPV1 (1:1000, Millipore Sigma, Cat# SAB5700857). The specificity of primary antibodies was evaluated by either western blot or pre-absorption assay by us or the manufactures. Immunostaining in the absence of primary or secondary antibody was assessed for background evaluation. After primary antibody overnight incubation and rinsing, fluorescence-conjugated species-specific secondary antibody (1:500, Molecular Probes, Eugene, OR) was applied for 2 h and at room temperature. The secondary antibodies used were donkey anti-rabbit (Cy3) (1:500, Jackson Immuno research, Cat# 711-165-152), donkey anti-goat 594 (1:500, Life Technologies, Cat# A11058), donkey anti-rabbit 488 (1:500, Life Technologies, A21206), and donkey anti-goat 488 (1:500, Life Technologies, Cat# A11055). Sections were finally applied a coverslip with Citifluor (Citifluor Ltd., London) mounting medium and viewed under a Zeiss fluorescent microscope.

### DRG single neuron Ca^2+^ imaging

Freshly obtained DRGs were subject to enzymatic disassociation in Gibco Dulbecco’s Modified Eagle Medium (DMEM) containing 2 mg/mL collagenase at 32 ℃ for 1 hour. After washing, DRG neurons were re-suspended into DMEM containing 10 % fetal bovine serum (FBS) and seeded into coated glass-bottom chamber for culture. Prior to experiments, DRG neurons were fasted for 2-4 hours. For Ca^2+^ imaging, GCaMP6f intensity was monitored and video recorded via a time-lapsing microscopic software in a rate of 1 frame/per second.

### Total RNA extraction, PCR, and quantitative (q)PCR

DRG tissues, spinal cord tissues, and cultured spinal cord astrocytes were subject to total RNA extraction using the RNAqueous™ Total RNA Isolation Kit (Thermo Fisher Scientific). Reverse transcription was performed using a cDNA synthesis kit High Capacity cDNA Reverse Transcription (Applied Biosystems). For conventional PCR (cPCR), products were assessed via agarose gel electrophoresis. For qPCR, SYBR Green was used as indicator on StepOnePlus™ Systems (Applied Biosystems). The expression level of β-actin in the same RNA extract sample was used as internal control for normalization of the target genes. The following were lists of primers used in cPCR: glial fibrillary acidic protein (GFAP): CCCTGGCTCGTGTGGATTT and GACCGATACCACTCCTCTGTC; Pgp9.5: GATGCTGAACAAAGTGTTGGC and GGAGTTTCCGATGGTCTGCTT; TrkB.T1 (*NTRK2.T1*): CCATTGCCCTCTGCTAACCA and GACAAAGTTGCTGCCTGGTG; TrkB (*NTRK2*): GCAAATCGCAGCAGGTATGG and TCAGAGCGAAGGACAGCAAG; beta (β)-actin (*ACTB*). The primers for qPCR were: tumor necrosis factor-alpha (TNFα): CAGGCGGTGCCTATGTCTC and CGATCACCCCGAAGTTCAGTAG; interleukin (IL)-6: CTGCAAGAGACTTCCATCCAG and AGTGGTATAGACAGGTCTGTTGG; *ACTB*: GGCTGTATTCCCCTCCATCG and CCAGTTGGTAACAATGCCATGT. Results of qPCR were calculated with ΔCt method and expressed as fold changes (2^-ΔΔCt^ fold).

### Induction of colitis

To induce colitis, we performed intracolonic installation of 2,4,6-trinitrobenzenesulfonic acid (TNBS, Sigma-Aldrich, St. Louis, MO) in a single dose (75 μL of 1.25 % solution prepared in 30 % EtOH) under anesthesia (2 % isoflurane). TNBS was delivered via a transanal polyethylene (PE)-50 catheter and infused to 3 cm inside from the anus. The animal was lifted by holding up the tail for 1 min after TNBS installation to avoid drug leakage from the anus. The same amount of EtOH was used as vehicle treatment and served as control.

### Somatic mechanical sensitivity

Hind paw responses to von Frey filament stimulation were assessed to indicate mechanical sensitivity. Briefly, mice were placed individually into plexiglas chambers sitting above a mesh stand (IITC Life Science Inc., CA). After acclimation to the environment for 30 min, a von Frey filament was applied perpendicularly to stimulate the plantar surface of the hind paw from underneath of the mesh floor when the animal had all four paws resting on the floor. The mice were each consecutively tested for a particular filament size, starting from the smallest, and this was repeated five rounds before moving on to a filament size of higher force. The tests were blinded to minimize bias. A behavior that was considered as positive included paw licking, shaking, or withdrawal (lifting) during or immediately after application. A painful response was defined by three positive responses out of the five stimulations for each filament size. A left shift of the response curve revealed mechanical hypersensitivity.

### Hot plate and tail flick assay

Hind paw thermal hyperalgesia was tested via a hot plate (52°C) assay. Specifically, a temperature-controlled metal plate coupled with a clear plexiglass cylinder (IITC Life Science Inc., CA) was used for testing. After the animal was placed into the cylinder, a time latency was measured between animal’s paw touching the plate till animal to show signs of and/or either lifting or licking of the hind paw. Each test was not exceeded beyond 30 secs to avoid tissue damage. For tail flick assay, the tail of the mice was submerged to hot water bath (52°C). The latency between the time of tail touching the water and the time the tail flicks out of the water was recorded.

### DRG explants culture

Freshly dissected DRG explant pairs from the same spinal segment were used for acute culture. Specifically, DRG explant was placed into a well of a 96-well plate containing DMEM. After DRGs were settled in cell culture incubator for 2-4 hours (fasting), one DRG from each DRG pair was treated with drugs and its contralateral DRG served as its control.

### Experimental Design and Statistical Analyses

#### Behavioral assessment

Behavioral assays were performed in a longitudinal manner and compared between groups. The comparison groups were tested on the same day in a blind fashion. When the animals received all behavioral tests on the same day, we started with mechanical sensitivity assay, followed by tail flick and then hot plate assay. Mechanical sensitivity tests and tail flick were examined in the morning block and the hot plate assay was performed in the afternoon so that animals can be rehabilitated.

#### Biochemical assay

For any comparison, animals were sacrificed on the same day and tissues were processed simultaneously in the same manner.

#### DRG neuron counting

The number of DRG neurons from a DRG section was counted in a blind fashion. Only those DRG neurons that showed visible nucleus were included. The area of the corresponding DRG section that contained neuronal soma was measured, avoiding any place that had nerve fibers. The result was calculated as the number of positive DRG neurons over the corresponding DRG section area as normalization. Data from multiple sections of a given DRG were pooled and averaged as one data point. Every third section in the serial cutting was stained for each specific antibody to avoid double counting.

#### Software and Statistical analysis

We used Imaging J, Zeiss ZEN pro, Nikon NIS-ELEMENTS-BR for data analysis. We used GraphPad Prism 9 for statistical analysis. The sample sizes were determined by power analysis to achieve an alpha of 0.05 with a power of 80 %. The number of animals used were described in the Result section in each experiment. One-way ANOVA, two-way ANOVA and *t* test were used. The results from each study were presented as mean ± SEM. Differences between means at a level of p≤0.05 were considered to be significant. Exact p values were included in each figure.

## RESULT

### Colitis increased the percentage of hind paw primary afferent neurons expressing Piezo2 and CGRP

We previously reported that hind paw mechanical sensitivity was enhanced on day 7 post induction of TNBS colitis in male mice [7]. To explain the hind paw behavioral outcomes, we examined the neurochemical profiles in retrograde dye FB labeled hind paw primary afferent neurons in L4 DRG on day 7 post colitis induction, emphasizing on Piezo2 and CGRP due to their roles in mechanical sensitivity and pain. In WT control mice, we found that a subset of Piezo2-expressing (Piezo2^+^) DRG neurons (**Fig 1A**, red cells) were hind paw primary afferent neurons that were labeled by FB (**Fig 1A**, blue cells). The percentage of FB-labeled L4 DRG neurons containing Piezo2 (**Fig 1A**, purple cells) was increased by colitis (**Fig 1B**, n=5 control, n=4 colitis, unpaired two-tailed *t* test, p=0.0001). Similarly, colitis increased the percentage of FB-labeled hind paw primary afferent neurons in L4 DRG expressing CGRP immunoreactivity (**Fig 1C**, red cells, **Fig 1D**, n=5 control, n=4 colitis, unpaired two-tailed *t* test, p=0.0001). Using two molecular markers we found that colitis-induced hind paw behavioral hypersensitivity could be explicated by the upregulation of Piezo2 and CGRP in hind paw primary afferent neurons.

**Fig 1.**
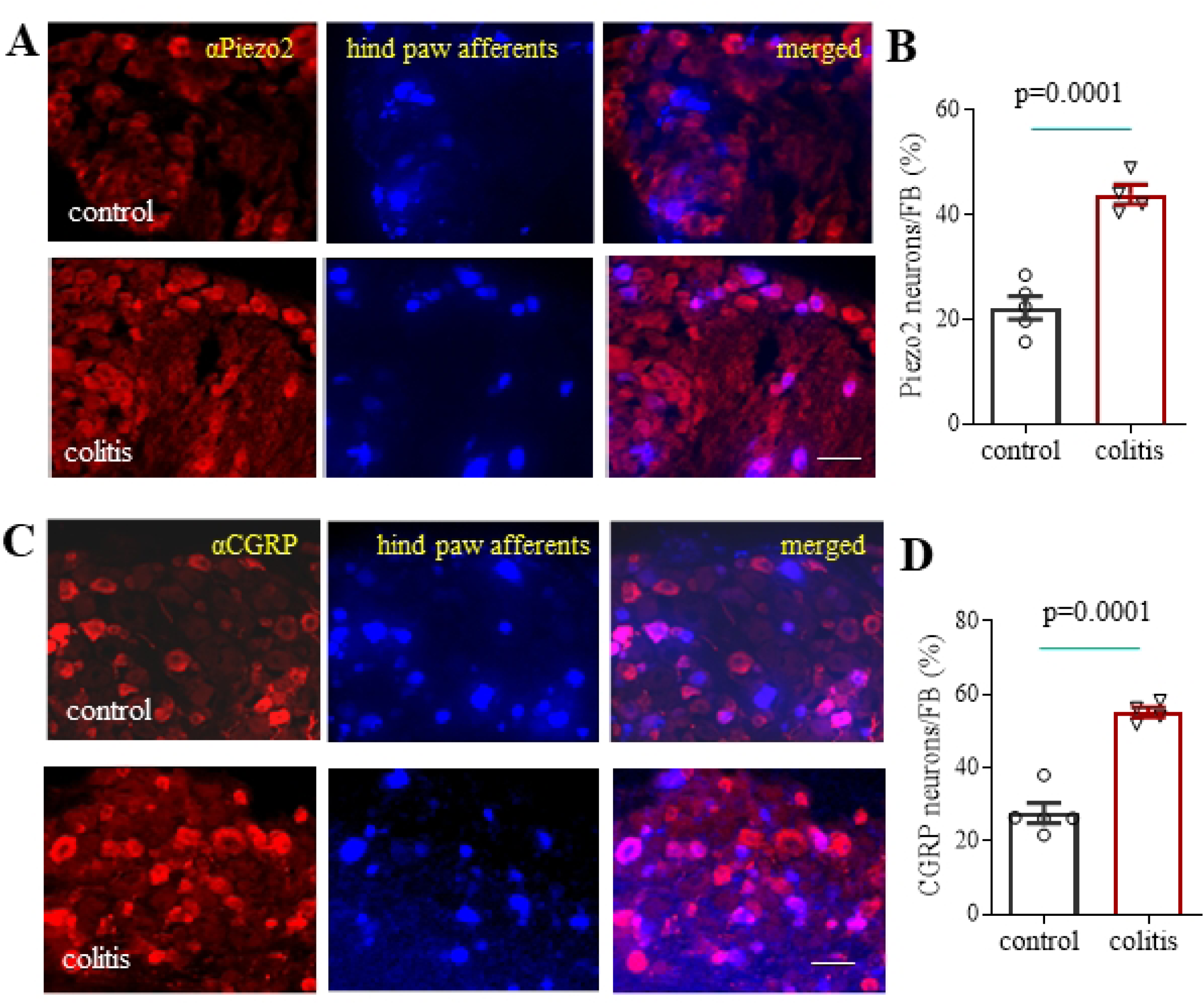
Colitis increases Piezo2 and CGRP expression in hind paw primary afferent neurons. (A) Expression of Piezo2 immunoreactivity (red cells) in hind paw primary afferent neurons labeled by Fast Blue (FB, blue cells) and co-expressing cells appear as purple after microphotograph merging. Bar = 50 μm. (B) Colitis increases the percentage of FB-labeled hind paw primary afferent neurons expressing Piezo2. (C) Expression of CGRP immunoreactivity (red cells) in hind paw primary afferent neurons labeled by FB (blue cells) and co-expressing cells appear as purple after microphotograph merging. Bar = 50 μm. (D) Colitis increases the percentage of FB-labeled hind paw primary afferent neurons expressing CGRP.

### Colitis-induced hind paw mechanical hypersensitivity as well as Piezo2 and CGRP upregulation in hind paw primary afferent neurons were mediated by endogenous TrkB.T1

Since BDNF is a potent neurotransmitter in pain facilitation, we examined the role of BDNF high affinity receptor TrkB.T1 in colitis-induced hind paw hypersensitivity by using TrkB.T1 knockout (KO) mice. We first compared the baseline hind paw mechanical sensitivity between wildtype (WT) and TrkB.T1 KO mice, which showed that TrkB.T1 KO did not affect hind paw baseline mechanical sensitivity when compared to counterpart WT mice (**Fig 2A**, n=10 WT, n=15 TrkB.T1 KO, two-way ANOVA, p values were presented in the Figure). We also compared hind paw thermal sensitivity at baseline between WT and TrkB.T1 KO mice which did not have significant differences (**Fig 2B**, n=17 WT, n=24 TrkB.T1 KO, unpaired two-tailed t test, p=0.7539). Similarly, a tail flick assay did not reveal differences of thermal hyperalgesia between WT and TrkB.T1 KO mice in naïve mice (**Fig 2C**, n=10 WT, n=11 TrkB.T1 KO, unpaired two-tailed t test, p=0.2043). These results suggested that TrkB.T1 KO did not produce compensatory mechanical and thermal behavioral abnormality in untreated states.

**Fig 2.**
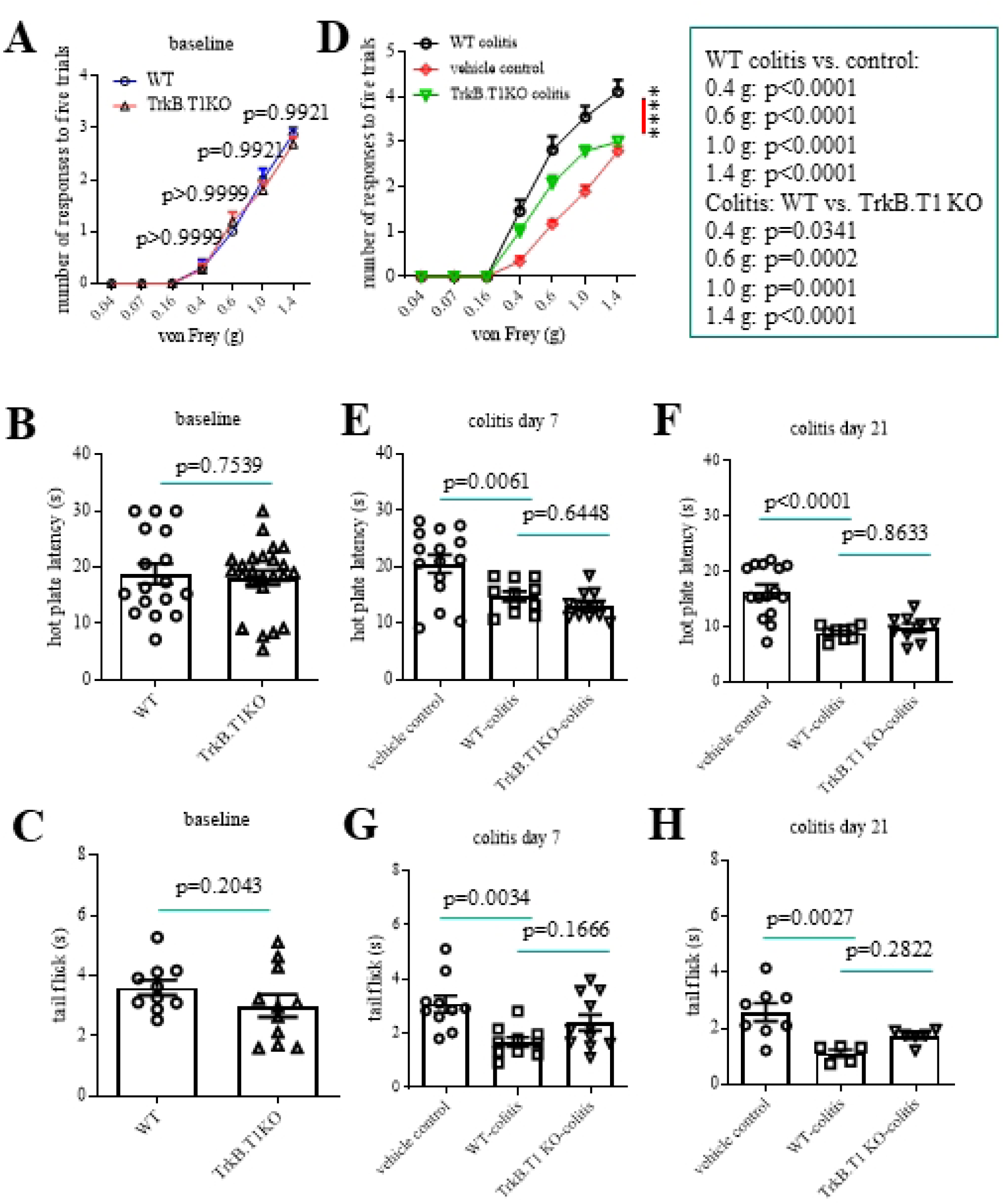
TrkB.T1 deletion attenuates colitis-induced hind paw mechanical hypersensitivity but not thermal hyperalgesia. (A) TrkB.T1 deletion does not change hind paw mechanical baseline sensitivity. (B) TrkB.T1 deletion does not change hind paw thermal baseline sensitivity. (C) TrkB.T1 deletion does not change baseline thermal hyperalgesia examined by tail flick. (D) TrkB.T1 deletion attenautes the development of hind paw mechanical hypersensitivity caused by colitis. (E-F) TrkB.T1 deletion has no effect on hind paw thermal hypersensitivity caused by colitis. (G-H) TrkB.T1 deletion has no effect on tail flick thermal hyperalgesia caused by colitis.

The assessments of hind paw mechanical sensitivity in TrkB.T1 KO mice with TNBS colitis (day 7) revealed that TrkB.T1 KO significantly reduced colitis-induced hind paw mechanical hypersensitivity when compared to WT colitic mice (**Fig 2D**, compare green line with black line) (Two-way ANOVA, n=18 vehicle control, n= 11 WT colitis, n=14 TrkB.T1 KO colitis, p values were presented in the Figure); this reduction occurred in response to noxious stimulation (bigger than 0.6 g von Frey filament weight), suggesting that endogenous TrkB.T1 participated in colitis-induced hind paw mechanical pain development. Notably, TrkB.T1 KO did not ameliorate colitis-induced hind paw thermal hypersensitivity examined on both day 7 (**Fig 2E**, one-way ANOVA, p values were indicated in the Figure, n=15 vehicle control, n=11 WT colitis, n=12 TrkB.T1 KO colitis) and day 21 (**Fig 2F**, one-way ANOVA, p values were indicated in the Figure, n=15 vehicle control, n=9 WT colitis, n=9 TrkB.T1 KO colitis), nor affected colitis-induced thermal hyperalgesia assessed by tail flick on both day 7 (**Fig 2G**, one-way ANOVA, p values were indicated in the Figure, n=10 vehicle control, n=10 WT colitis, n=11 TrkB.T1 KO colitis) and day 21 (**Fig 2H**, one-way ANOVA, p values were indicated in the Figure, n=8 vehicle control, n=5 WT colitis, n=6 TrkB.T1 KO colitis).

Since colitis-induced hind paw hypersensitivity was accompanied with the increased number of hind paw primary afferent neurons expressing Piezo2 and CGRP (**Fig 1**), we examined the role of TrkB.T1 in these molecular profiles. We compared the results between WT colitic (day 7) and TrkB.T1 KO colitic (day 7) mice and found that the percentage of FB-labeled hind paw primary afferent neurons (**Fig 3A**, blue cells) containing Piezo2 (**Fig 3A**, purple cells) was significantly lower in TrkB.T1 KO colitic mice when compared to WT colitic mice (**Fig 3B**, n=4 WT colitis, n=4 TrkB.T1 KO colitis, unpaired two-tailed *t* test, p=0.0022). Similarly, the percentage of FB-labeled hind paw primary afferent neurons (**Fig 3C**, blue cells) containing CGRP (**Fig 3C**, teal cells) was also significantly lower in TrkB.T1 KO colitic mice when compared to WT colitic mice (**Fig 3D**, n=5 WT colitis, n=5 TrkB.T1 KO colitis, unpaired two-tailed *t* test, p=0.0045). These results suggested that TrkB.T1-mediated hind paw mechanical hypersensitivity was manifested by an increase in the percentage of hind paw primary afferent neurons expressing Piezo2 or CGRP.

**Fig 3.**
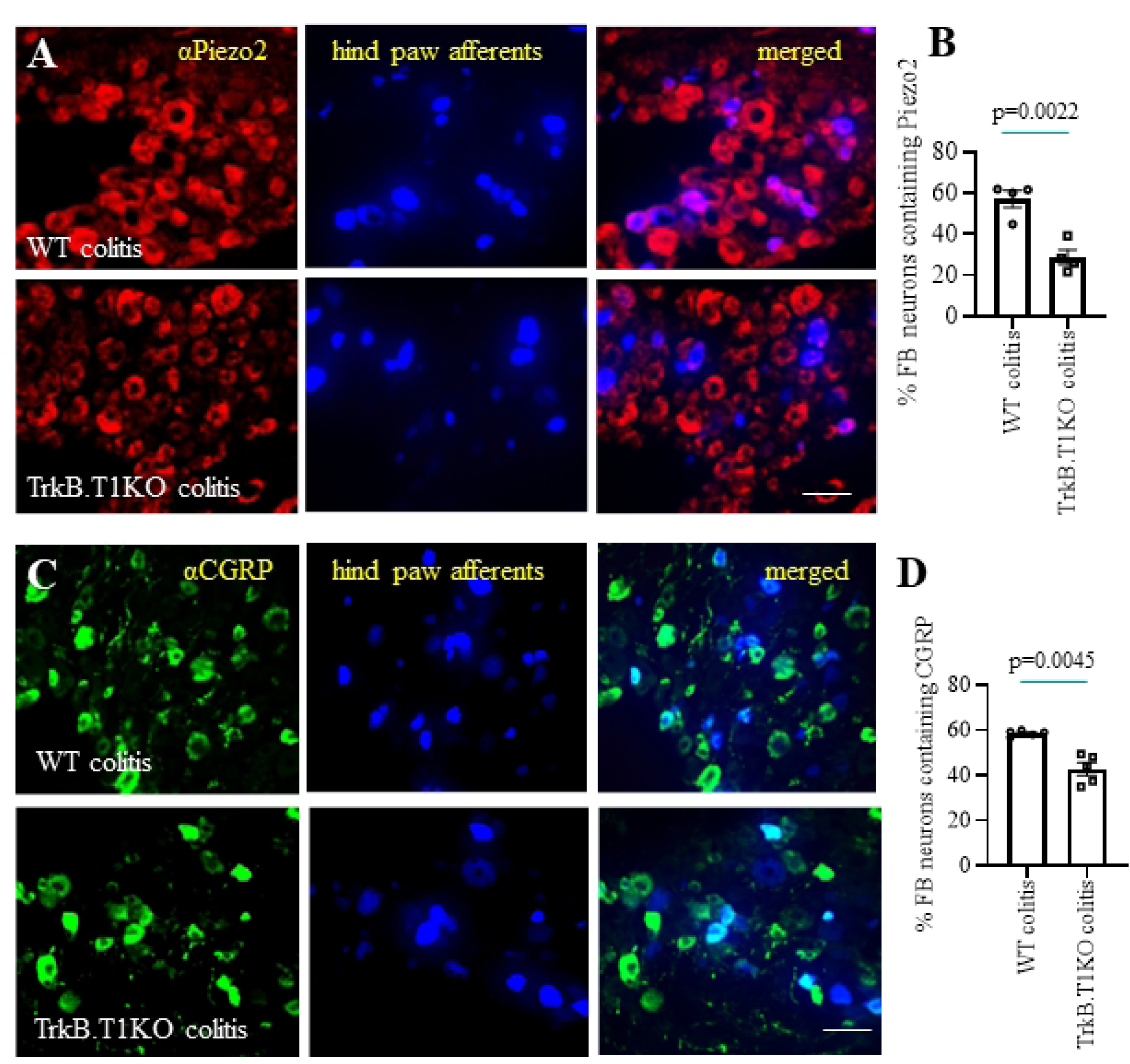
Effects of TrkB.T1 KO on colitis-induced Piezo2 and CGRP upregulation in hind paw primary afferent neurons. (A) Expression of Piezo2 immunoreactivity (red cells) in hind paw primary afferent neurons labeled by FB (blue cells) and co-expressing cells appear as purple after microphotograph merging. Bar = 50 μm. (B) TrkB.T1 KO reduces the percentage of hind paw primary afferent neurons expressing Piezo2 in colitis. (C) Expression of CGRP immunoreactivity (green cells) in hind paw primary afferent neurons labeled by FB (blue cells) and co-expressing cells appear as teal after microphotograph merging. Bar = 50 μm. (D) TrkB.T1 KO reduces the percentage of hind paw primary afferent neurons expressing CGRP in colitis.

### TrkB.T1 mediated TNFα upregulation in the spinal cord in colitis to regulate Piezo2 and CGRP expression in DRG

Neuroinflammation plays a critical role in the spinal central sensitization that mediates spinal trans-segmental crosstalk between colonic and hind paw primary afferent neurons (**Fig 4A**). TNF-α and IL-6 are key pro-inflammatory cytokines in the generation of neuroinflammation in the spinal cord, contributing to chronic pain development [45–47]. We measured the mRNA levels of TNFα (**Fig 4B**) and IL-6 (**Fig 4C**) in the spinal cord in colitis and found that both cytokines were upregulated. However, colitis-induced TNFα upregulation was attenuated by TrkB.T1 KO (**Fig 4B**, one-way ANOVA, p values were indicated in the Figure, n=6 for vehicle control, n=6 for WT colitis, n=4 for TrkB.T1 KO colitis) while the IL-6 level was not affected by TrkB.T1 KO (**Fig 4C**, one-way ANOVA, p values were indicated in the Figure, n=6 for vehicle control, n=6 for WT colitis, n=4 for TrkB.T1 KO colitis). Colitis also increased the expression level of TrkB.T1 in the spinal cord of WT mice (**Fig 4D**, n=6 control, n=6 colitis, unpaired two-tailed *t* test, p=0.0006). We examined the cell type-specific mRNA expression of TrkB.T1 and found that TrkB.T1 was expressed by the spinal cord astrocytes that did not contain the full-length TrkB mRNA, using the whole DRG mRNA extracts (contained all molecules examined) as positive control (**Fig 4E**, full gel images were attached as supplement materials). These results suggested that colitis increased TNFα expression in the spinal cord, presumably from astrocytes, via TrkB.T1-mediated cellular events (**Fig 4F**).

**Fig 4.**
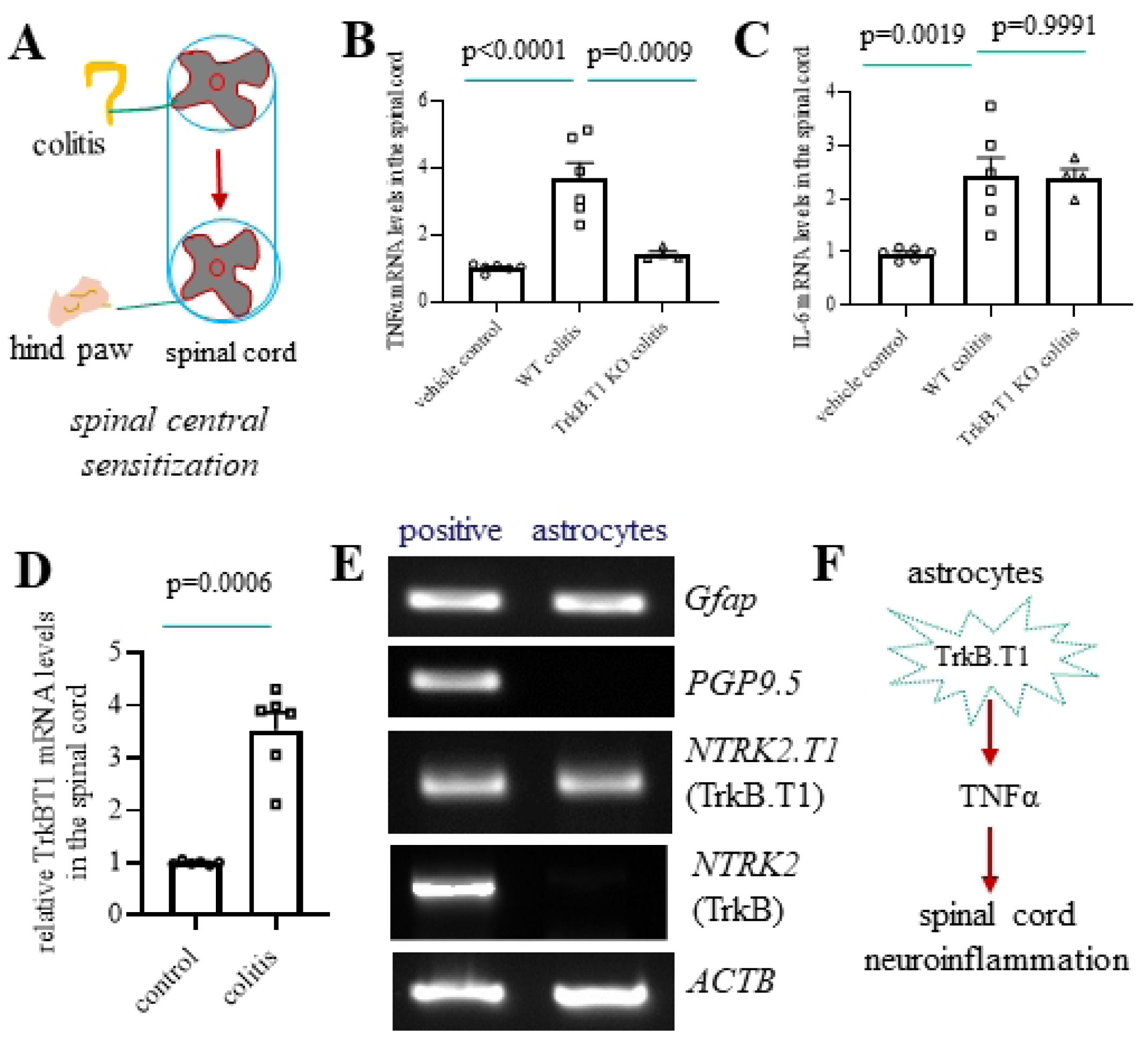
TrkB.T1 mediates TNFα production in the spinal cord in colitis. (A) Diagram shows an involvement of spinal central sensitization in colon-to-hind paw cross-organ sensitization. (B) TrkB.T1 mediates colitis-induced TNFα upregulation in the spinal cord. (C) TrkB.T1 does not have a role in colitis-induced IL-6 upregulation in the spinal cord. (D) The TrkB.T1 mRNA level is increased in the spinal cord by colitis. (E) TrkB.T1 but not TrkB is expressed by spinal cord astrocytes. (F) Diagram summarizes TrkB.T1 in astrocytes in mediation of TNFα production.

Since TrkB.T1 was involved in colitis-induced Piezo2 and CGRP upregulation in hind paw primary afferent neurons (**Fig 3**), we wondered whether TNFα as a downstream mediator of TrkB.T1 was able to regulate Piezo2 and CGRP expression in DRG. To do so, we first tested the responsiveness of DRG neurons to TNFα stimulation via Ca^2+^ imaging, in which we were able to customize the concentration of TNFα used to exert effective stimulation (0.1 ng/mL, 1 ng/mL, 10 ng/mL). We cultured DRG neurons from mice containing the organic Ca^2+^ indicator GCaMP6f and found that TNFα (1 ng/mL) stimulation increased the intracellular Ca^2+^ levels in a subset of DRG neurons (**Fig 5A**, **5B**). The subsequent application of capsaicin (0.1 µM) following TNFα treatment revealed that a subset of DRG neurons responsive to TNFα stimulation were either capsaicin-insensitive (**Fig 5B)** or capsaicin-sensitive (**Fig 5C**), with the final step of treatment with KCl (100 mM) to validate that these DRG neurons were still alive following TNFα and capsaicin treatment (**Fig 5C-D**). These results suggested that only a part of DRG neurons responded to TNFα treatment and some of them were nociceptors. After the TNFα treatment of DRG explants culture, we found that TNFα increased the number of DRG neurons expressing Piezo2 (**Fig 5E, 5F,** n=5, paired two-tailed *t* test, p=0.0033) or CGRP (**Fig 5G, 5H,** n=5, paired two-tailed *t* test, p=0.0015). Since the PI3K/Akt pathway was regulated by TNFα in DRG neurons and participated in somatic pain [48, 49], we also examined whether the PI3K/Akt pathway participated in TNFα-induced Piezo2 and CGRP upregulation in DRG. Our results showed that TNFα-induced Piezo2 upregulation was inhibited by the PI3K/Akt inhibitor LY294002 (LY, 10 μM; **Fig 5I**, n=4, paired two-tailed *t* test, p=0.0094)), while TNFα-induced CGRP upregulation in DRG neurons was not affected by LY 294002 (**Fig 5J**, n=4 for each group, paired two-tailed *t* test, p=0.1908). These results suggested multiple signaling pathways were likely involved in colitis-induced hind paw hypersensitivity.

**Fig 5.**
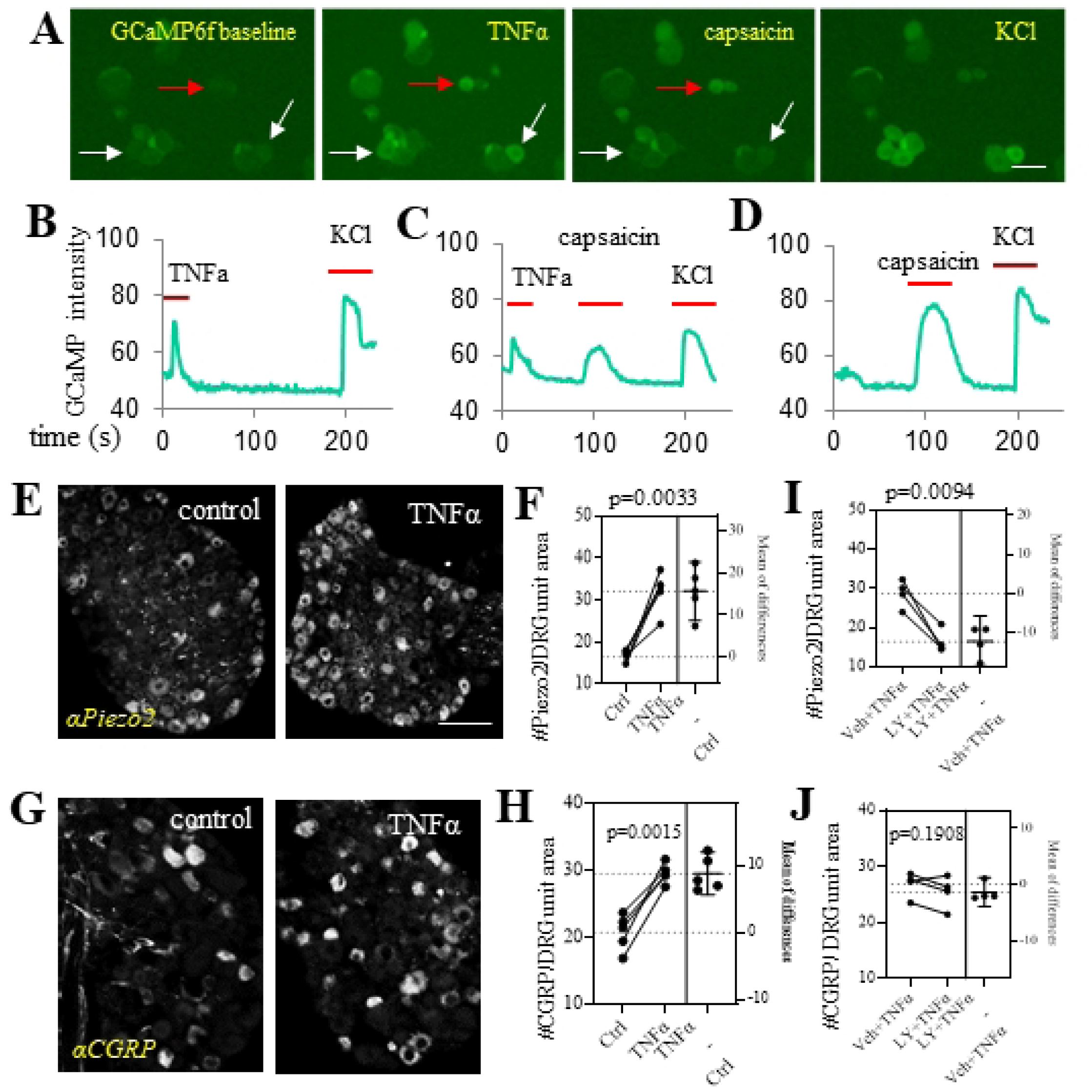
TNFα sensitizes DRG neurons and induces Piezo2 and CGRP expression in DRG differentially mediated by the PI3K/Akt pathway. (A) Single DRG neuron calcium imaging in response to TNFα stimulation followed by capsaicin treatment and KCl live neuron validation. Bar = 100 μm. (B) Calcium transients in DRG neurons responsive to TNFα but not capsaicin treatment. (C) Calcium transients in DRG neurons responsive to both TNFα and capsaicin treatments. (D) Calcium transients in DRG neurons responsive to capsaicin but not TNFα treatment. (E) Piezo2 immunoreactivity in DRG neurons in response to TNFα treatment. Bar = 100 μm. (F) TNFα treatment increases the number of DRG neurons expressing Piezo2. (G) CGRP immunoreactivity in DRG neurons in response to TNFα treatment. Bar = 100 μm. (H) TNFα treatment increases the number of DRG neurons expressing CGRP. (I) Inhibition of the PI3K/Akt pathway reduces TNFα-facilitated Piezo2 expression in DRG neurons. (J) Inhibition of the PI3K/Akt pathway does not have significant effects on TNFα-facilitated CGRP expression in DRG neurons.

### TrkB.T1 mediated the activation of Akt in Piezo2-expressing hind paw primary afferent neurons in colitis

Since the PI3K/Akt pathway mediated Piezo2 upregulation in DRG neurons in culture, we examined the phosphorylation level (active form) of Akt (p-Akt) in L4 DRG in vivo and its association with Piezo2 in colitis. We used Piezo2;mCitrine mice that were generated by breeding Piezo2-Cre mice with R26-LSL-Gi-DREADD mice so that Piezo2-expressing (Piezo2^+^) cells could be visualized by mCitrine yellow fluorescent protein (YFP). We performed immunostaining to detect p-Akt in DRG neurons. We found that colitis increased the number of DRG neurons expressing p-Akt (**Fig 6A**, red cells; **Fig 6B**, n=8, unpaired two-tailed *t* test, p=0.0081), or Piezo2 (YFP) (**Fig 6A**, green cells; **Fig 6C**; n=3, unpaired two-tailed *t* test, p=0.0073) as well as co-expressing p-Akt and Piezo2 (**Fig 6A**, yellow cells indicated by arrows; **Fig 6D**, n=3, unpaired two-tailed *t* test, p=0.0018). Consistent to Piezo2, the upregulation of p-Akt in L4 DRG by colitis was also attenuated by TrkB.T1 KO (**Fig 6E**, n=4, unpaired two-tailed *t* test, p=0.0025; **Fig 6F**, red cells). A subset of FB-labeled hind paw primary afferent neurons (**Fig 6F**, blue cells) contained p-Akt (**Fig 6F**, purple cells) in colitis, which was markedly reduced by TrkB.T1 KO (**Fig 6G**, n=4 for each group, unpaired two-tailed *t* test, p=0.0177). These results suggested that colitis-induced TrkB.T1-mediated Piezo2 upregulation in hind paw primary afferent neurons involved an elevation of the PI3K/Akt activation.

**Fig 6.**
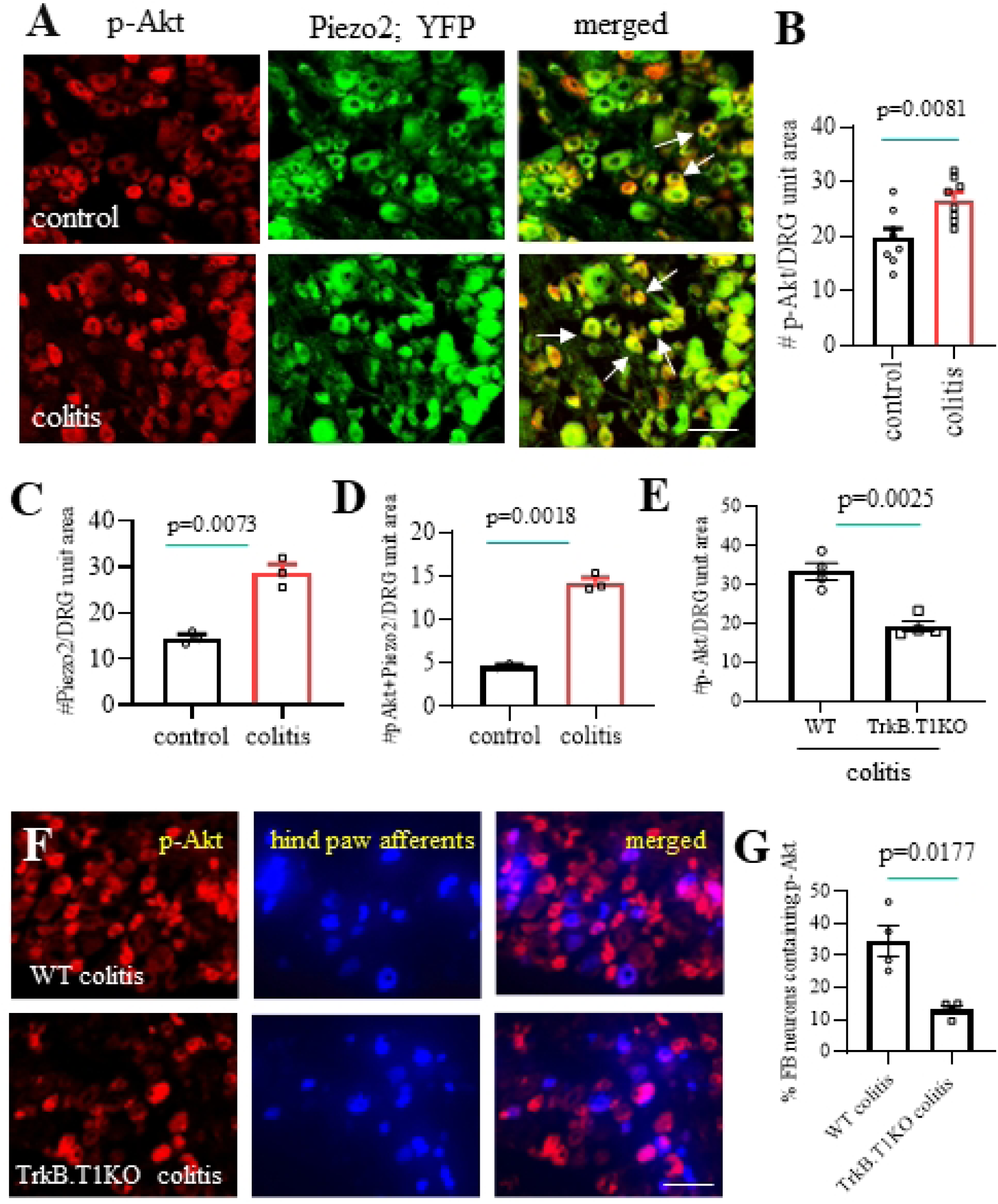
Colitis-induced Piezo2 upregulation in hind paw primary afferent neurons is associated with Akt activation. (A) Co-expression of p-Akt (red cells) and Piezo2;YFP (green cells) in L4 DRG neurons, and the co-expressing cells appear yellow after microphotograph merging (indicated by arrows). Bar = 50 μm. (B) Colitis increases the number of L4 DRG neurons expressing p-Akt. (C) Colitis increases the number of L4 DRG neurons expressing Piezo2 (YFP). (D) Colitis increases the number of L4 DRG neurons co-expressing Piezo2 and p-Akt. (E) TrkB.T1 KO reduces the number of L4 DRG neurons expressing p-Akt in colitis. (F) Expression of p-Akt immunoreactivity (red cells) in hind paw primary afferent neurons labeled by FB (blue cells) and co-expressing cells appear as purple after microphotograph merging. Bar = 50 μm. (G) TrkB.T1 KO reduces the percentage of hind paw primary afferent neurons expressing p-Akt in colitis.

### Colitis-induced CGRP upregulation in hind paw primary afferent neurons was not associated with p-Akt but with p-CREB

Assessment of CGRP and p-Akt co-expression in L4 DRG in colitis showed that these two molecules were scarcely overlapped (**Fig 7A**; CGRP: green cells; p-Akt: red cells), which explained the non-involvement of the PI3K/Akt pathway in CGRP upregulation in DRG in vitro (**Fig 5**). Since TNFα was able to evoke Ca2+ transients and increase CGRP expression in DRG neurons (**Fig 5**), we examined the co-expression of CGRP with the phosphorylation form of cAMP response element-binding protein (p-CREB). The activation/phosphorylation of CREB was dependent of the intracellular Ca^2+^ levels and was essential for CGRP transcription by binding to CGRP promoter. We found that a significant co-expression of CGRP (**Fig 7B**, green cells) with p-CREB (**Fig 7B**, red nuclear stain) in L4 DRG neurons occurred in colitis (**Fig 7B**, merged microphotograph). We next characterized the association of p-CREB and CGRP in FB-labeled hind paw primary afferent neurons in colitis and examined the effects of TrkB.T1 KO on their expression (**Fig 7C-E**). We found that a subpopulation of p-CREB/CGRP co-expressing neurons were FB-labeled hind paw (Hpaw) primary afferent neurons (**Fig 7C**, indicated by arrows). We also found that TrkB.T1 KO in colitis reduced the percentage of hind paw primary afferent neurons expressing p-CREB when compared to WT colitis (**Fig 7D**, n=4, unpaired two-tailed *t* test, p=0.0001). However, the percentage of CGRP^+^ DRG neurons expressing p-CREB was similar in WT colitis and TrkB.T1 colitis (**Fig 7E**, n=4, unpaired two-tailed *t* test, p=0.1013), suggesting a parallel regulation of p-CREB and CGRP expression in L4 DRG by TrkB.T1-mediated molecular and cellular events in colitis.

**Fig 7.**
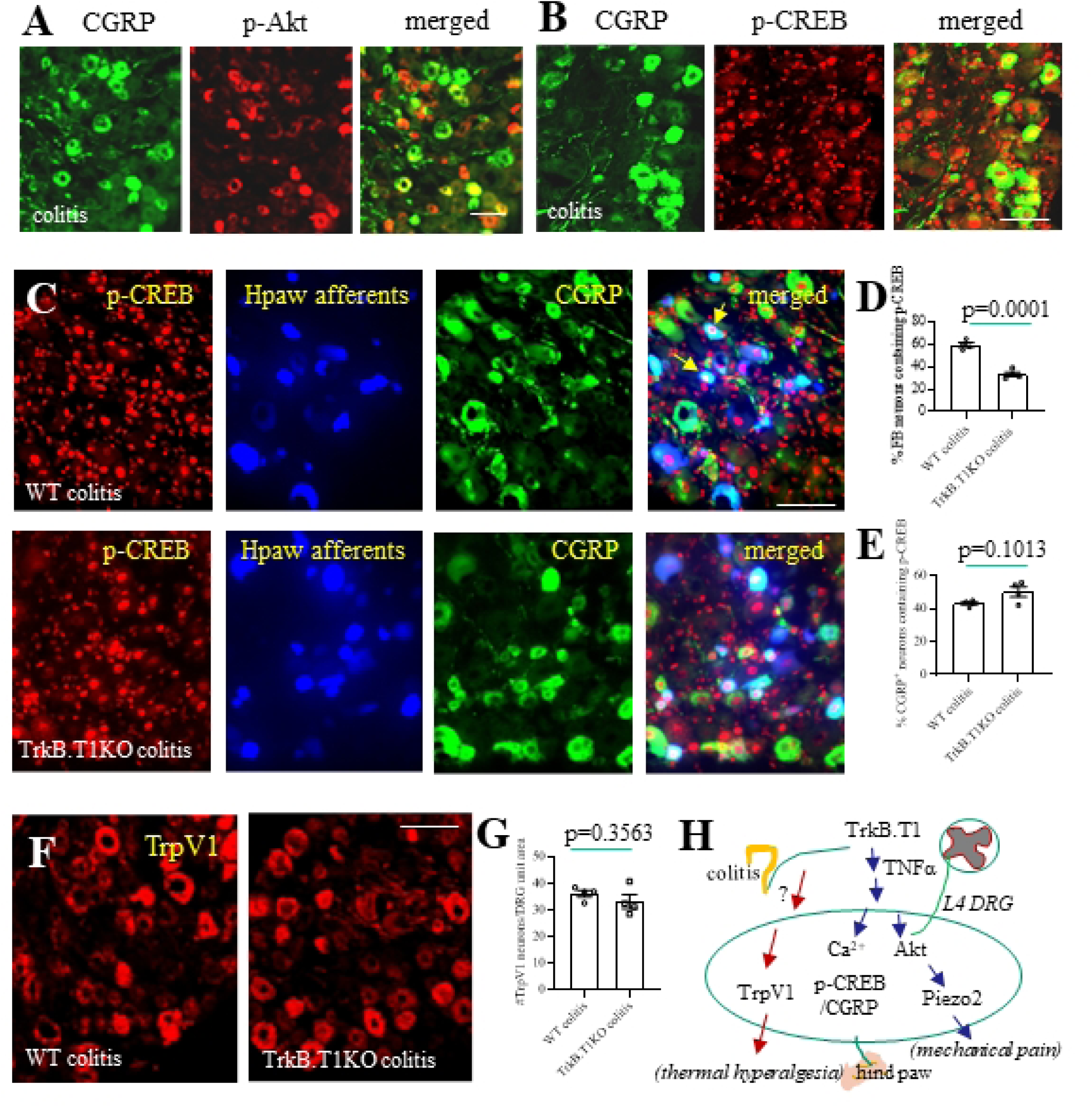
Effects of TrkB.T1 KO on p-CREB-CGRP axis and TrpV1 expression in L4 DRG in colitis. (A) Distribution of CGRP-positive (green cells) and p-Akt positive (red cells) L4 DRG neurons in colitis showing a few neurons containing both molecules (yellow cells). Bar = 50 μm. (B) Distribution of CGRP-positive (green cells) and p-CREB positive (red nuclear stain) L4 DRG neurons in colitis showing CGRP-positive neurons contain p-CREB. Bar = 50 μm. (C) Distribution of p-CREB immunoreactivity (red nuclear stain) in hind paw (Hpaw) primary afferent neurons (blue cells) and/or CGRP-positive neurons (green cells) of the same DRG section shows co-expression (indicated by arrows). Bar = 50 μm. (D) TrkB.T1 KO reduces the percentage of hind paw primary afferent neurons expressing p-CREB in colitis. (E) TrkB.T1 KO has no effects on the percentage of CGRP-positive L4 DRG neurons expressing p-CREB. (F) TrpV1 immunoreactivity (red cells) in L4 DRG in colitis. Bar = 50 μm. (G) TrkB.T1 KO has no effects on the number of L4 DRG neurons expressing TrpV1 in colitis. (H) Signal transduction explaining the role of TrkB.T1 in colitis-induced neurochemical changes in L4 DRG underlying colon-to-hind paw mechanical and thermal hypersensitivity.

### TrkB.T1 did not mediate colitis-induced TrpV1 expression in L4 DRG

Since that some DRG neurons were responsive to both TNFα and capsaicin (**Fig 5C**), we compared TrpV1 expression in L4 DRG between wildtype colitic and TrkB.T1 KO colitic mice (**Fig 7F**, red cells). We found that TrkB.T1 KO did not have significant effects on the number of DRG neurons expressing TrpV1 in L4 DRG in colitis (**Fig 7G**, n=4, unpaired two-tailed *t* test, p=0.3563). These results suggested that TNFα could activate but might not increase the number of DRG neurons expressing TrpV1 in L4 DRG neurons in colitis thus TrkB.T1-mediated TNFα action did not contribute to hind paw thermal hyperalgesia (**Fig 2**). Taken together, colitis-induced hind paw hypersensitivity necessitates the increase in the number of DRG neurons to express specific molecules such as Piezo2 to transduce mechanical pain and TrkB.T1-TNFα axis in the spinal cord, likely from astrocytes, is associated with the referred mechanical but not thermal pain development (**Fig 7H**).

## DISCUSSION

The present study demonstrates that hind paw mechanical pain as a result of colitis involves TrkB.T1-mediated TNFα upregulation in the spinal cord as well as TrkB.T1-mediated Piezo2 and CGRP upregulation in hind paw primary afferent neurons in L4 DRG. Specifically, TrkB.T1 deletion attenuates colitis-induced hind paw mechanical hypersensitivity but not thermal hyperalgesia, suggesting a unique role of TrkB.T1 in the development of distinct somatic pain modalities. In the spinal cord that links colonic afferents and hind paw afferents to convey colon-to-hind paw sensory signals, TrkB.T1 KO reduces colitis-induced upregulation of proinflammatory cytokine TNFα but not IL-6, in turn, TNFα activates DRG neurons and increases the expression level of Piezo2 and CGRP in DRG. Mechanistically, the PI3K/Akt pathway is associated with Piezo2 but not CGRP upregulation in DRG by colitis or TNFα; p-CREB is associated with CGRP upregulation in hind paw primary afferent neurons in colitis. TrkB.T1 does not participate in colitis-induced hind paw thermal hyperalgesia nor TrpV1 expression in L4 DRG. These studies at the behavioral and molecular levels suggest a role of TrkB.T1 in mediating colitis-induced hind paw referred mechanical but not thermal hypersensitivity that is associated with TNFα upregulation in the spinal cord.

A health problem exists in IBS patients who also experience somatic pain in the limbs. A clinical study comparing between 42 IBS patients and 29 control subjects shows that both diarrhea-predominant (n=27) and constipation-predominant (n=15) IBS patients display hypersensitivity to ischemic arm pain while healthy participants do not [50]. IBS Patients (10 women, 2 men) also demonstrate cutaneous allodynia/hyperalgesia to thermal pain applied to the hand and foot when compared to control (13 women, 4 men), with cutaneous hyperalgesia more pronounced in the foot than hand [51]. A meta-analysis of 4 cross-sectional studies and one cohort study involving 86,438 individuals shows that patients with IBS have a nearly three-fold increased odds of restless legs syndrome when compared with controls [52]. It is also common (up to one-third) in patients with IBD to have joint pain and problem in the skin, eye, and mouth [53]. IBD-associated joint pain (arthralgia) is usually symmetric with no direct inflammation/damage to the joint (referred pain), which is different from Arthritis that only affects the inflamed joint [53]. These widespread extraintestinal manifestations greatly worsen the quality of life and necessitates a better understanding of the underlying mechanisms to guide for treatment. Using rat and mouse models, we and others show that colitis induced by localized intracolonic installation of TNBS triggers mechanical and thermal hypersensitivity examined in the hind foot [4, 5, 8], proving that this model system can be used to examine the molecular pathways that govern the colon-to-somatic organ cross-sensitization in order to understand the mechanisms underlying the referred pain.

Primary afferent neurons in DRG are organized in strict orders to ensure precise sensation of the environmental cues such as touch, prick, distention, heat, chemical stimulation, and others. Each distinct organ is innervated by sensory neuron nerve fibers stemming from distinct spinal segments. For example, the colonic afferent neurons are housed in thoracolumbar and lumbar sacral segments [10], while the lumbar L4 DRG innervate the hind limb. Interestingly, in response to colonic inflammation not only the colonic afferent neurons are sensitized but the neurons in L4 DRG are also activated showing as increased number of hind paw primary afferent neurons expressing Piezo2 and CGRP. Visceral irritation from intracolonic capsaicin instillation also increases the percentage of L4 DRG neurons having an elevated Ca^2+^ response to hind paw pinch [8]. It remains unclear for the pathways to convey signals from colonic afferent neurons to hind paw afferent neurons. Here we show that TrkB.T1 that is primarily expressed by astrocytic glia mediates colitis-induced hind paw mechanical hypersensitivity as well as the expression of Piezo2 and CGRP in hind paw primary afferent neurons. Mice lacking TrkB.T1 demonstrate reduced hind paw mechanical hypersensitivity after colitis induction and also reduced expression of Piezo2 and CGRP in hind paw primary afferent neurons. However, colitis-induced hind paw thermal hyperalgesia remains unaffected by TrkB.T1 deletion. These results suggested that antagonism of TrkB.T1 may be effective to treat referred mechanical pain but it won’t work on thermal pain.

Both of the expression levels of TNFα and IL-6 are increased in the spinal cord by colitis. In colitic mice lacking TrkB.T1, the upregulation of TNFα in the spinal cord by colitis is attenuated while the upregulation of IL-6 is not affected, suggesting that TrkB.T1-mediated a specific pathway involving TNFα in the colon-to-hind paw cross-organ sensitization. Indeed, TNFα treatment sensitizes nociceptive neurons and increases the expression levels of Piezo2 and CGRP in DRG. IL-6 is likely produced by microglia that do not express TrkB.T1 [27, 54, 55], explaining that colitis-induced IL-6 upregulation in the spinal cord is not reduced in mice lacking TrkB.T1. A previous study using optogenetic approaches shows that chronic activation of spinal astrocytes evokes hind paw mechanical pain and thermal hyperalgesia [56]. Comparing to the role of TrkB.T1 in astrocytes that only mediates mechanical hypersensitivity but not thermal hyperalgesia that are both induced by colitis, it suggests a diverse population of astrocytes in the spinal cord among which non-TrkB.T1-expressing subpopulations may participate in thermal hyperalgesia, which is beyond the scope of the current study. Another note is on TNFα. TNF antagonist etanercept (1 mg, i.p., every third day) attenuates spinal nerve ligation-induced mechanical allodynia, however, with its action in DRG but not in the spinal cord [57]. In the spinal cord, TNF-α evokes a drastic increase in spontaneous excitatory postsynaptic current frequency in lamina II neurons which participates in thermal hyperalgesia [58]. The discrepancy between TrkB.T1-mediated behavioral changes and TNFα action could be due to that TNFα is also produced by other immune cells in the spinal cord such as microglia, similar as IL-6 [47, 59, 60]. Taken together, colon-to-hind paw cross-organ sensitization involves multiple signaling pathways and TrkB.T1-mediated TNFα production is just one of them to facilitate the referred mechanical pain.

Although colitis induces both Piezo2 and CGRP upregulation in L4 DRG, distinct signaling pathways are involved. Colitis-induced Piezo2 upregulation is associated with the activation of PI3K/Akt pathway, while colitis-induced CGRP upregulation is associated with the activation of transcription factor CREB. When using TNFα to upregulate Piezo2 and CGRP, we find that TNFα-induced Piezo2 upregulation can be attenuated by the PI3K/Akt inhibitor, however, TNFα-induced CGRP upregulation is not affected by inhibition of the PI3K/Akt pathway. These findings further suggest that multiple signaling networks participate, in parallel, in the development of colitis-induced somatic mechanical hypersensitivity. The findings that TrkB.T1 KO does not affect IL-6 expression in the spinal cord and TrpV1 expression in L4 DRG neurons as well as hind paw thermal hyperalgesia suggested distinct mechanisms underlying colitis-induced somatic mechanical vs thermal pain.

### Summary and limitations

The current study provides previously uncharacterized mechanisms and signaling pathways that underlie interorgan hetero-segmental sensory crosstalk. Our findings that TrkB.T1 mediates neuroinflammation in the spinal cord to trigger somatic pain in colitis provide information to guide the development of therapeutic strategies to treat referred pain by targeting neuroinflammatory signals in the spinal cord. Our findings that reveal multiple complicated signaling networks in colon-to-hind paw cross-organ sensitization suggest that this health problem is very complex and necessitates in-depth studies to identify effective therapeutic targets. The limitation of this study is that TrkB.T1 KO deletes all TrkB.T1 in the body. Although TrkB.T1 is expressed in the spinal astrocytes and mediates TNFα upregulation in the spinal cord in colitis, the cell-type specific role of TrkB.T1 necessitates examinations in future studies. If astrocytes are indeed involved in the process, whether it is caused by astrocytic cytokine trans-segmental diffusion or due to gap junction-mediated intercellular Ca^2+^ waves within astrocyte networks as seen in the brain [61] necessitates further investigation.

## Data Availability

all data is included in the manuscript. Source data is attached.

## Acknowledgements

We thank Dr. Jose M Eltit from Virginia Commonwealth University help with calcium imaging set-up, Dr. Lino Tessarollo from NIH to provide TrkB.T1 knockout mice, and Dr. Divya Sharma to assist in the qPCR experiment.

